# Ninety-nine independent genetic loci influencing general cognitive function include genes associated with brain health and structure (N = 280,360)

**DOI:** 10.1101/176511

**Authors:** Gail Davies, Max Lam, Sarah E Harris, Joey W Trampush, Michelle Luciano, W David Hill, Saskia P Hagenaars, Stuart J Ritchie, Riccardo E Marioni, Chloe Fawns-Ritchie, David CM Liewald, Judith A Okely, Ari V Ahola-Olli, Catriona LK Barnes, Lars Bertram, Joshua C Bis, Katherine E Burdick, Andrea Christoforou, Pamela DeRosse, Srdjan Djurovic, Thomas Espeseth, Stella Giakoumaki, Sudheer Giddaluru, Daniel E Gustavson, Caroline Hayward, Edith Hofer, M Arfan Ikram, Robert Karlsson, Emma Knowles, Jari Lahti, Markus Leber, Shuo Li, Karen A Mather, Ingrid Melle, Derek Morris, Christopher Oldmeadow, Teemu Palviainen, Antony Payton, Raha Pazoki, Katja Petrovic, Chandra A Reynolds, Muralidharan Sargurupremraj, Markus Scholz, Jennifer A Smith, Albert V Smith, Natalie Terzikhan, Anbu Thalamuthu, Stella Trompet, Sven J van der Lee, Erin B Ware, B Gwen Windham, Margaret J Wright, Jingyun Yang, Jin Yu, David Ames, Najaf Amin, Philippe Amouyel, Ole A Andreassen, Nicola J Armstrong, Amelia A Assareh, John R Attia, Deborah Attix, Dimitrios Avramopoulos, David A Bennett, Anne C Böhmer, Patricia A Boyle, Henry Brodaty, Harry Campbell, Tyrone D Cannon, Elizabeth T Cirulli, Eliza Congdon, Emily Drabant Conley, Janie Corley, Simon R Cox, Anders M Dale, Abbas Dehghan, Danielle Dick, Dwight Dickinson, Johan G Eriksson, Evangelos Evangelou, Jessica D Faul, Ian Ford, Nelson A Freimer, He Gao, Ina Giegling, Nathan A Gillespie, Scott D Gordon, Rebecca F Gottesman, Michael E Griswold, Vilmundur Gudnason, Tamara B Harris, Annette M Hartmann, Alex Hatzimanolis, Gerardo Heiss, Elizabeth G Holliday, Peter K Joshi, Mika Kähönen, Sharon LR Kardia, Ida Karlsson, Luca Kleineidam, David S Knopman, Nicole A Kochan, Bettina Konte, John B Kwok, Stephanie Le Hellard, Teresa Lee, Terho Lehtimäki, Shu-Chen Li, Tian Liu, Marisa Koini, Edythe London, Will T Longstreth, Oscar L Lopez, Anu Loukola, Tobias Luck, Astri J Lundervold, Anders Lundquist, Leo-Pekka Lyytikäinen, Nicholas G Martin, Grant W Montgomery, Alison D Murray, Anna C Need, Raymond Noordam, Lars Nyberg, William Ollier, Goran Papenberg, Alison Pattie, Ozren Polasek, Russell A Poldrack, Bruce M Psaty, Simone Reppermund, Steffi G Riedel-Heller, Richard J Rose, Jerome I Rotter, Panos Roussos, Suvi P Rovio, Yasaman Saba, Fred W Sabb, Perminder S Sachdev, Claudia Satizabal, Matthias Schmid, Rodney J Scott, Matthew A Scult, Jeannette Simino, P Eline Slagboom, Nikolaos Smyrnis, Aïcha Soumaré, Nikos C Stefanis, David J Stott, Richard E Straub, Kjetil Sundet, Adele M Taylor, Kent D Taylor, Ioanna Tzoulaki, Christophe Tzourio, André Uitterlinden, Veronique Vitart, Aristotle N Voineskos, Jaakko Kaprio, Michael Wagner, Holger Wagner, Leonie Weinhold, K Hoyan Wen, Elisabeth Widen, Qiong Yang, Wei Zhao, Hieab HH Adams, Dan E Arking, Robert M Bilder, Panos Bitsios, Eric Boerwinkle, Ornit Chiba-Falek, Aiden Corvin, Philip L De Jager, Stéphanie Debette, Gary Donohoe, Paul Elliott, Annette L Fitzpatrick, Michael Gill, David C Glahn, Sara Hägg, Narelle K Hansell, Ahmad R Hariri, M Kamran Ikram, J. Wouter Jukema, Eero Vuoksimaa, Matthew C Keller, William S Kremen, Lenore Launer, Ulman Lindenberger, Aarno Palotie, Nancy L Pedersen, Neil Pendleton, David J Porteous, Katri Räikkönen, Olli T Raitakari, Alfredo Ramirez, Ivar Reinvang, Igor Rudan, Dan Rujescu, Reinhold Schmidt, Helena Schmidt, Peter W Schofield, Peter R Schofield, John M Starr, Vidar M Steen, Julian N Trollor, Steven T Turner, Cornelia M Van Duijn, Arno Villringer, Daniel R Weinberger, David R Weir, James F Wilson, Anil Malhotra, Andrew M McIntosh, Catharine R Gale, Sudha Seshadri, Thomas H. Mosley, Jan Bressler, Todd Lencz, Ian J Deary

## Abstract

General cognitive function is a prominent human trait associated with many important life outcomes^1,2^, including longevity^3^. The substantial heritability of general cognitive function is known to be polygenic, but it has had little explication in terms of the contributing genetic variants^4,5,6^. Here, we combined cognitive and genetic data from the CHARGE and COGENT consortia, and UK Biobank (total N=280,360; age range = 16 to 102). We found 9,714 genome-wide significant SNPs (*P*<5 x 10^−8^) in 99 independent loci. Most showed clear evidence of functional importance. Among many novel genes associated with general cognitive function were *SGCZ*, *ATXN1*, *MAPT*, *AUTS2*, and *P2RY6*. Within the novel genetic loci were variants associated with neurodegenerative disorders, neurodevelopmental disorders, physical and psychiatric illnesses, brain structure, and BMI. Gene-based analyses found 536 genes significantly associated with general cognitive function; many were highly expressed in the brain, and associated with neurogenesis and dendrite gene sets. Genetic association results predicted up to 4% of general cognitive function variance in independent samples. There was significant genetic overlap between general cognitive function and information processing speed, as well as many health variables including longevity.

---

Since its discovery in 1904^7^, hundreds of studies have replicated the finding that around 40% of the variance in people’s test scores on a diverse battery of cognitive tests can be accounted for by a single general factor^8^. General cognitive function is peerless among human psychological traits in terms of its empirical support and importance for life outcomes^1,2^. Individual differences in general cognitive function show phenotypic and genetic stability across most of the life course^9-11^. Twin studies find that general cognitive function has a heritability of more than 50% from adolescence through adulthood to older age^4,12,13^. SNP-based estimates of heritability for general cognitive function are about 20-30%^5^. To date, little of this substantial heritability has been explained; only a few relevant genetic loci have been discovered (**Table 1** and **Fig. 1**). Like other highly polygenic traits, a limitation on uncovering relevant genetic loci is sample size^14^; to date, there have been fewer than 100,000 individuals in studies of general cognitive function^5,6^.

**Table 1.**
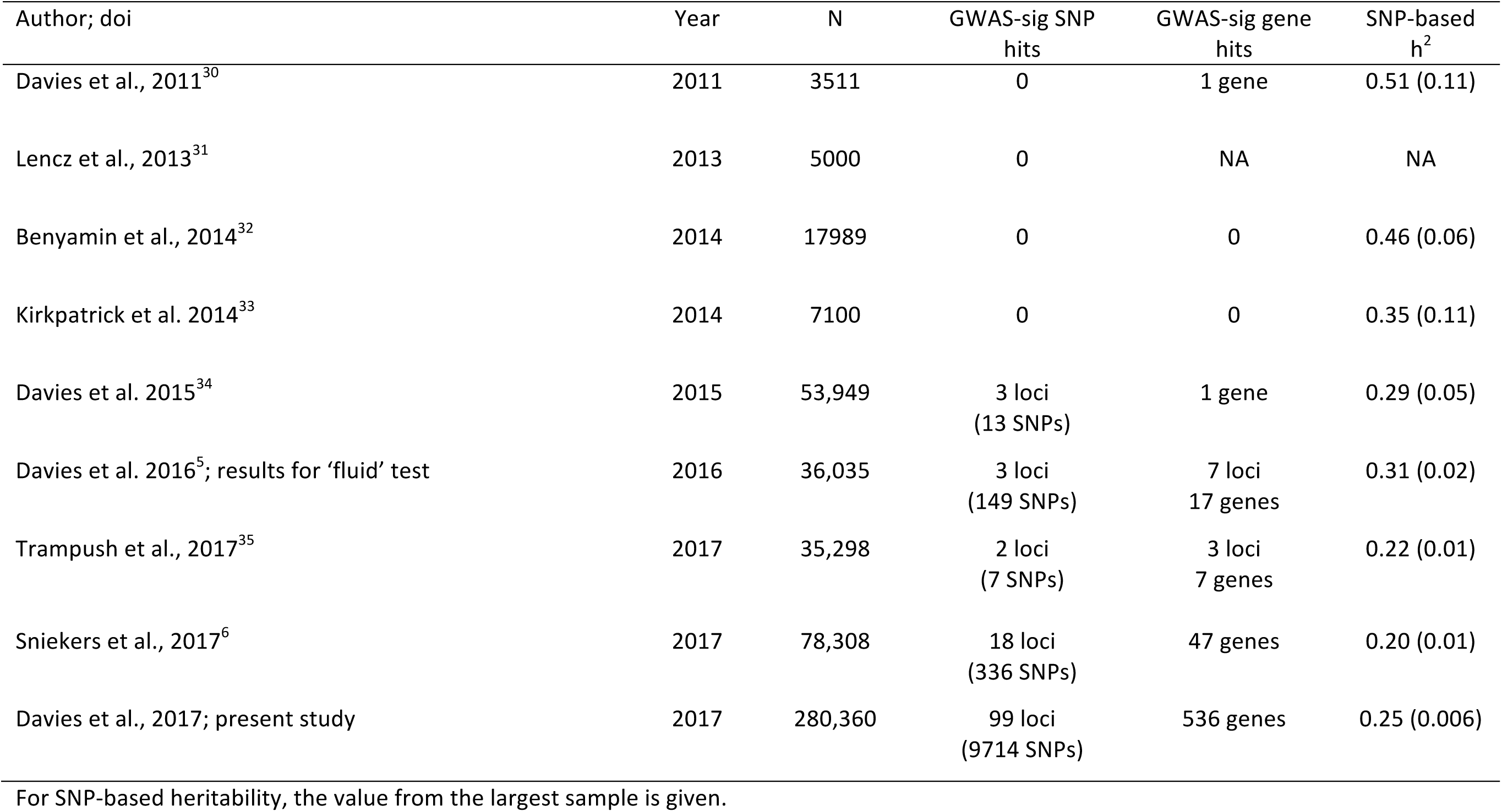
Details of GWA studies of general cognitive function to date, including the present study

**Figure 1.**
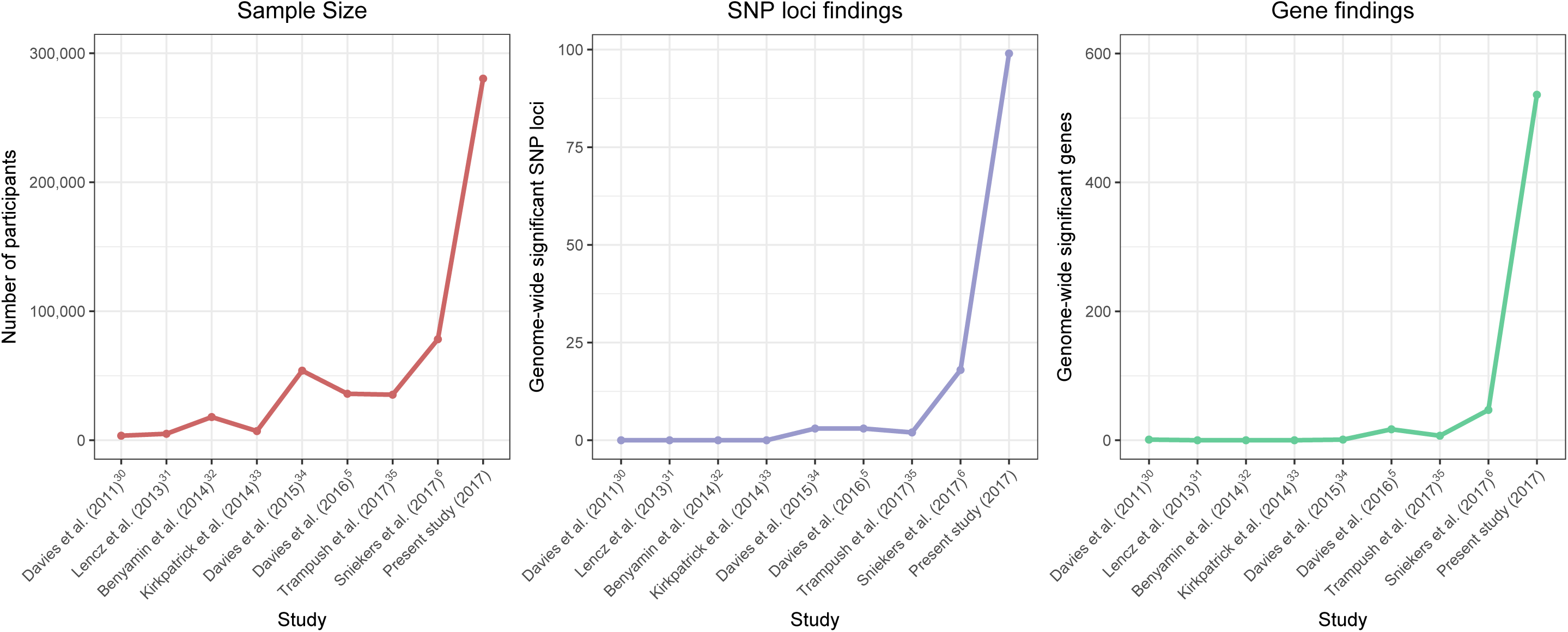
Summary of molecular genetic association studies with general cognitive function to date.

General cognitive function, unlike height for example, is not measured the same way in all samples. Here, this was mitigated by applying a consistent method of extracting a general cognitive function component from cognitive test data in the cohorts of the CHARGE and COGENT consortia; all individuals were of European ancestry (**Supplementary Materials**). Cohorts’ participants were required to have scores from at least three cognitive tests, each of which tested a different cognitive domain. Each cohort applied the same data reduction technique (principal component analysis) to extract a general cognitive component. Scores from the first unrotated principal component were used as the general cognitive function phenotype. Using a general cognitive function phenotype in a genetically informative design is supported by the observation that the well-established positive manifold of cognitive tests is best presented by a highly heritable higher-order latent general cognitive function phenotype that mediates genetic and environmental covariances among cognitive tests^4,8,13^. The psychometric characteristics of the general cognitive component from each cohort in the CHARGE consortium are shown in **Supplementary Materials**. In order to address the fact that different cohorts had applied different cognitive tests, we previously showed that two general cognitive function components extracted from different sets of cognitive tests on the same participants correlate highly^5^. The cognitive test from the large UK Biobank sample was the so-called ‘fluid’ test, a 13-item test of verbal-numerical reasoning, which has a high genetic correlation with general cognitive function^15^. With the CHARGE and COGENT samples’ general cognitive function scores and UK Biobank’s verbal-numerical reasoning scores (in two samples: assessment centre-tested, and online-tested), there were 280,360 participants included in the present report’s genome-wide association study (GWAS) analysis. We performed two post-GWAS meta-analyses separately: first, on the CHARGE and COGENT cohorts; and, second, on UK Biobank’s two samples. Prior to running the subsequent meta-analysis of CHARGE-COGENT with UK Biobank, the genetic correlation, calculated using linkage disequilibrium score (LDSC) regression, was estimated at 0.82 (SE=0.02), indicating very substantial overlap between the genetic variants influencing general cognitive function in these two groups. We performed an inverse-variance weighted meta-analysis of CHARGE-COGENT and UK Biobank.

Genome-wide results for general cognitive function showed 9,714 significant (*P* < 5 × 10^−8^) SNP associations, and 17,563 at a suggestive level (1 × 10^−5^ > *P* > 5 × 10^−8^); see **Fig. 2a** and **Supplementary Tables 3 and 4**. There were 120 independent lead SNPs identified by FUnctional MApping and annotation of genetic associations (FUMA)^16^. A comparison of these lead SNPs with results from the largest previous GWAS of cognitive function^6^ and educational attainment^17^—which included a subsample of individuals contributing to the present study—confirmed that 4 and 12 of these, respectively, were genome-wide significant in the previous studies (Supplementary Table 14). Five SNPs in the present study were completely novel (i.e., *P* > .05 in these previous studies): rs7010173 (chromosome 8; intronic variant in *SGCZ*); rs179994 (chromosome 6; intronic variant in *ATXN1*), rs8065165 (chromosome 17; intronic variant 2KB upstream of *MAPT*); rs2007481 (chromosome 7; intronic variant in *AUTS2*); and rs188236525 (chromosome 11; intronic variant 2KB upstream of *P2RY6*). The 120 lead SNPs were distributed within 99 loci across all autosomal chromosomes. Using the GWAS catalog (https://www.ebi.ac.uk/gwas/)” www.ebi.ac.uk/gwas/) to look up each locus, only 12 of these loci had been reported previously for other GWA studies of cognitive function or educational attainment (novel loci are indicated in **Supplementary Table 16**). Therefore, our study uncovered 87 novel independent loci associated with cognitive function. Of the five completely novel loci, two of these are in/near interesting candidate genes: *MAPT* gene mutations are associated with neurodegenerative disorders such as Alzheimer’s disease and frontotemporal dementia^18^; and *AUTS2* is a candidate gene for neurological disorders such as autism spectrum disorder, intellectual disability, developmental delay^19^, and for alcohol consumption^20,21^. These general cognitive function-associated genes also showed significant gene associations in the gene-based tests (except for *P2RY6*); see **Supplementary Table 7 and Fig. 2b** for the results for 536 genes that the present study finds to be significantly associated with general cognitive function.

**Figure 2.**
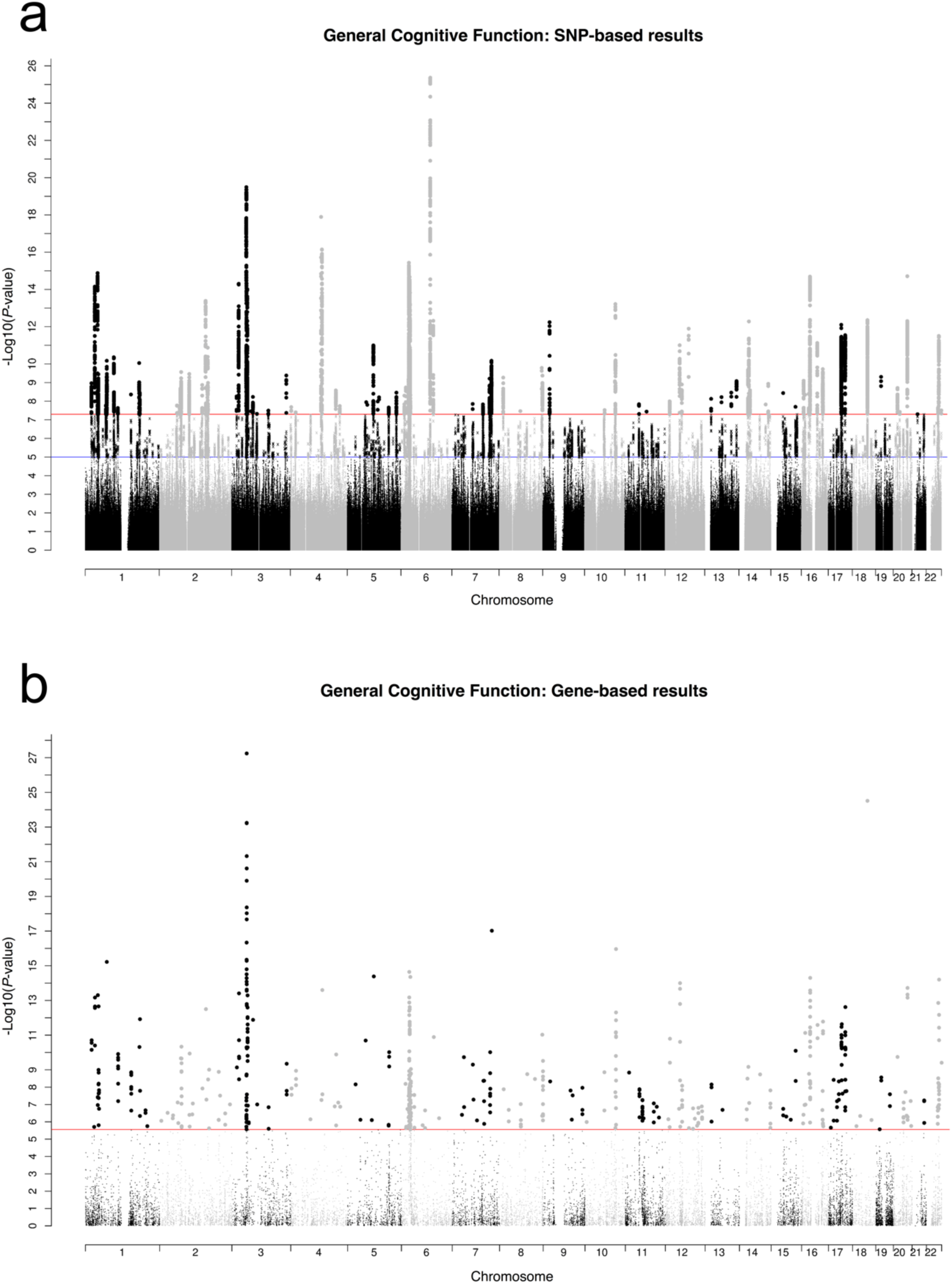
SNP-based (a) and gene-based (b) association results for general cognitive function in 280,360 individuals. The red line indicates the threshold for genome-wide significance: *P* < 5 × 10^−8^ for (a), *P* < 2.75 × 10^−6^ for (b); the blue line in (a) indicates the threshold for suggestive significance: *P* < 1 × 10^−5^.

For the 120 lead SNPs, a summary of previous SNP associations is listed in **Supplementary Table 15**. They have been associated with many physical (e.g., BMI, height, weight), medical (e.g., lung cancer, Crohn’s disease, blood pressure), and psychiatric (e.g., bipolar disorder, schizophrenia, autism) traits, as well as with cognitive function and educational attainment (12 loci). Of the novel SNP associations, we highlight previous associations with autism/ADHD (3 loci), bipolar disorder/schizophrenia (14 loci), and infant head circumference/intracranial volume/subcortical brain region volumes (2 loci).

We sought to identify lead and tagged SNPs within the 99 significant genomic risk loci associated with general cognitive function that are potentially functional, using FUMA^16^ (**Supplementary Table 16**). See online methods for further details. Seventy-nine of the genomic risk loci contained at least one SNP with a Combined Annotation Dependent Depletion (CADD) score > 12.37, indicating that they are likely to be deleterious SNPs. Sixty-five of the genomic risk loci contained at least one SNP with a RegulomeDB score < 3, indicating that they are likely to be involved in gene regulation. Ninety-seven of the loci contained at least one SNP with a minimum 15-core chromatin state score of < 8, indicating that they are located in an open chromatin state consistent with the SNP being in a regulatory region. Sixty-eight of the loci contained at least one eQTL. Of interest, rs1135840 in *CYP2D6* (P=1.42 × 10^−11^) is a non-synonymous SNP (Ser486Thr), that has previously been associated with the metabolism of several commonly-used drugs^22^.

MAGMA gene-set analysis identified two significant gene sets associated with general cognitive function: neurogenesis (*P* = 1.1 × 10^−7^) and dendrite (*P* = 1.6 × 10^−6^) (**Supplementary Table 18; see Online Methods**). Identification of these gene sets is consistent with genes associated with cognitive function regulating the generation of cells within the nervous system, including the formation of neuronal dendrites. MAGMA gene-property analysis indicated that genes expressed in all brain regions—except the brain spinal cord and cervical c1—and genes expressed in the pituitary share a higher level of association with general cognitive function than genes not expressed in the brain or pituitary (**Fig. 3** and **Supplementary Tables 20 and 21**). The most significant enrichments were for genes expressed in the cerebellum and the brain’s cortex.

**Figure 3.**
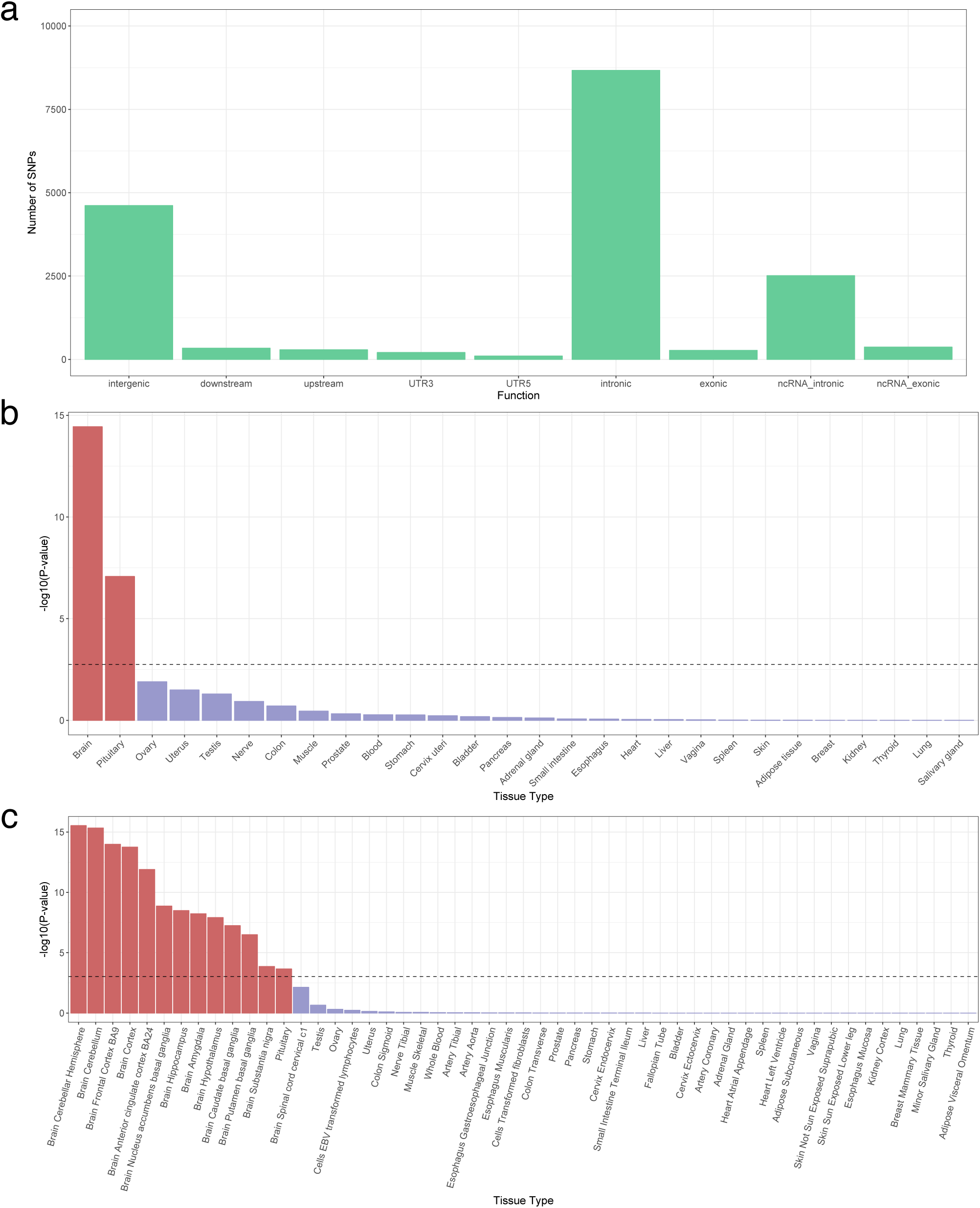
Functional analyses of general cognitive function association results, lead SNPs, and all SNPs in LD with lead SNPs. Functional consequences of SNPs on genes (a) indicated by functional annotation assigned by ANNOVAR. MAGMA gene-property analysis results; results are shown for average expression of 30 general tissue types (b) and 53 specific tissue types (c). The dotted line indicates the Bonferroni-corrected α level.

We estimated the proportion of variance explained by all common SNPs in four of the largest individual samples, using univariate GCTA-GREML analyses (see Online Methods): English Longitudinal Study of Ageing (ELSA: N = 6,661, *h*^2^ = 0.12, SE = 0.06), Understanding Society (N = 7,841, *h*^2^ = 0.17, SE = 0.04), UK Biobank Assessment Centre (N = 86,010, *h*^2^ = 0.25, SE = 0.006), and Generation Scotland (N = 6,507, *h*^2^ = 0.20, SE = 0.05^23^) (**Table 2**). Genetic correlations for general cognitive function amongst these cohorts, estimated using bivariate GCTA-GREML, ranged from *r*_*g*_ = 0.88 to 1.0 (**Table 2**). There were slight differences in the test questions and the testing environment for the UK Biobank’s ‘fluid’ (verbal-numerical reasoning) test in the assessment centre versus the online version. Therefore, we investigated the genetic contribution to the stability of individual differences in people’s verbal-numerical reasoning using a bivariate GCTA-GREML analysis, including only those individuals who completed the test on both occasions (mean time gap = 4.93 years). We found a significant perfect genetic correlation of *r*_*g*_ = 1.0 (SE = 0.02).

**Table 2.** Genetic correlations and heritability estimates of a general cognitive function component in three United Kingdom cohorts

| Cohort | ELSA | US | GS |
| --- | --- | --- | --- |
| ELSA | 0.12 (0.06) |  |  |
| US | 1.0 (0.33) | 0.17 (0.04) |  |
| GS | 1.0 (0.38) | 0.88 (0.24) | 0.20 (0.05) |

Below the diagonal, genetic correlations (standard error) of general cognitive function amongst three cohorts are shown: English Longitudinal Study of Ageing (ELSA); Generation Scotland (GS); and Understanding Society (US). SNP-based heritability (standard error) estimates appear on the diagonal.

We tested how well the genetic results from our CHARGE-COGENT-UK Biobank general cognitive function GWAS analysis accounted for cognitive test score variance in independent samples. We re-ran the GWAS analysis excluding three of the larger cohorts: ELSA, Generation Scotland, and Understanding Society. These new GWAS summary results were used to create polygenic profile scores in the three cohorts. The polygenic profile score for general cognitive function explained 2.37% of the variance in ELSA (*β* = 0.16, SE = 0.01, *P* = 1.40 × 10^−46^), 3.96% in Generation Scotland (*β* = 0.21, SE = 0.01, *P* = 3.87 × 10^−72^), and 4.00% in Understanding Society (*β* = 0.21, SE = 0.01, *P* = 1.31 × 10^−81^). Full results for all five thresholds are shown in **Supplementary Table 11**.

Using the CHARGE-COGENT-UK Biobank GWAS results, we tested the genetic correlations between general cognitive function and 25 health traits. Sixteen of the 25 health traits were significantly genetically correlated with general cognitive function (**Supplementary Table 12**). Novel genetic correlations were identified between general cognitive function and ADHD (*r*_*g*_ = -0.36, SE = 0.03, *P* = 3.91 × 10^−32^), bipolar disorder (*r*_*g*_ = -0.09, SE = 0.04, *P* = 0.008), major depression (*r*_*g*_ = -0.30, SE = 0.05, *P* = 4.13 × 10^−12^), and longevity (*r*_*g*_ = 0.15, SE = 0.06, *P* = 0.014).

We explored the genetic foundations of reaction time and its genetic association with general cognitive function. Reaction time is an elementary cognitive task that assesses a person’s information processing speed. It is both phenotypically and genetically correlated with general cognitive function, and accounts for some of its association with health^24,25^. We note the limitation that the UK Biobank’s reaction time variable is based on only four trials per participant. Full results and methods are in Supplementary materials. There were 330,069 individuals in the UK Biobank sample with both reaction time and genetic data. GWAS results for reaction time uncovered 2,022 significant SNPs in 42 independent genomic regions; 122 of these SNPs overlapped with general cognitive function, with 76 having a consistent direction of effect (sign test *P* = 0.008) (**Supplementary Table 9**). These genomic loci showed clear evidence of functionality (**Supplementary Table 17**). Using gene-based GWA, 191 genes attained statistical significance (**Supplementary Table 8**), 28 of which overlapped with general cognitive function (**Supplementary Table 10**). Gene-sets constructed using expression data indicated a role for genes expressed in the brain (**Supplementary Tables 22 and 23; Supplementary Fig. 3**). There was a genetic correlation (*r*_*g*_) of 0.227 (*P* = 4.33 × 10^−27^) between reaction time and general cognitive function. The polygenic score for reaction time explained 0.43% of the general cognitive function variance in ELSA (*P* = 1.42 × 10^−9^), 0.56 % in Generation Scotland (*P* = 2.49 × 10^−11^), and 0.26% in Understanding Society (*P* = 1.50 × 10^−6^).

People with higher general cognitive function are broadly healthier^26, 27^; here, we find overlap between genetic loci for general cognitive function and a number of physical health traits. These shared genetic associations may reflect a causal path from cognitive function to disease, cognitive consequences of disease, or pleiotropy^28^. For psychiatric illness, conditions like schizophrenia (and, to a lesser extent, bipolar disorder) are characterised by cognitive impairments^29^, and thus reverse causality (i.e. from cognitive function to disease) is less likely. In terms of localising more proximal structural and functional causes of variation in cognitive function, researchers could prioritise the genetic loci uncovered here that overlap with brain-related measures.

General cognitive function has prominence and pervasiveness in the human life course, and it is important to understand the environmental and genetic origins of its variation in the population^4^. The unveiling here of many new genetic loci, genes, and genetic pathways that contribute to its heritability (**Supplementary Tables 3, 7 and 18**; **Fig. 2**)—which it shares, as we find here, with many health outcomes, longevity, brain structure, and processing speed—provides a foundation for exploring the mechanisms that bring about and sustain cognitive efficiency through life.

## Acknowledgments

This research was conducted in The University of Edinburgh Centre for Cognitive Ageing and Cognitive Epidemiology, funded by the Biotechnology and Biological Sciences Research Council and Medical Research Council (MR/K026992/1). This research was conducted using the UK Biobank Resource (Application Nos. 10279 and 4844). Cohort-specific acknowledgements are in the Supplementary Materials.

## Contributions

GD and IJD drafted the manuscript with contributions from M Luciano, SEH, WDH, SJR, SPH, CF-R, and JO. GD, JWT and M Lam performed quality control of the CHARGE-COGENT data. IJD designed and overviewed the cognitive psychometric analyses in the CHARGE cohorts. GD and REM performed quality control of UK Biobank data. GD, JWT and M Lam analysed the data. SEH, WDH, SPH and M Luciano performed/assisted with downstream analysis. GD and IJD co-ordinated the CHARGE and UK Biobank work, and their integration with COGENT; TL, JWT and M Lam co-ordinated the COGENT work. All authors supplied phenotype data, genotype data, and GWA results, and commented on and approved the manuscript.

## Online Methods

### Participants and Cognitive Phenotypes

This study includes 280,360 individuals of European ancestry from 57 population-based cohorts brought together by the Cohorts for Heart and Aging Research in Genomic Epidemiology (CHARGE), the Cognitive Genomics Consortium (COGENT) consortia, and UK Biobank. All individuals were aged between 16 and 102 years. Exclusion criteria included clinical stroke (including self-reported stroke) or prevalent dementia.

For each of the CHARGE and COGENT cohorts, a general cognitive function component phenotype was constructed from a number of cognitive tasks. Each cohort was required to have tasks that tested at least three different cognitive domains. Principal component analysis was applied to the cognitive test scores to derive a measure of general cognitive function. Principal component analyses results for the CHARGE cohorts were checked by one author (IJD) to establish the presence of a single component. Scores on the first unrotated component were used as the cognitive phenotype (general cognitive function). UK Biobank participants were asked 13 multiple-choice questions that assessed verbal and numerical reasoning (VNR: UK Biobank calls this the ‘fluid’ test). The score was the number of questions answered correctly in two minutes. Two samples of UK Biobank participants with verbal-numerical reasoning scores were used in the current analysis. The first sample (VNR Assessment Centre) consists of UK Biobank participants who completed the verbal-numerical reasoning test at baseline in assessment centres (n = 107,586). The second sample (VNR Web-Based) consists of participants who did not complete the verbal-numerical reasoning test at baseline but did complete this test during the web-based cognitive assessment online (n = 54,021). Details of the cognitive phenotypes for all cohorts can be found in Supplementary Information Section 2.

At the baseline UK Biobank assessment, 496,790 participants completed the reaction time test. Details of the test can be found in Supplementary Information Section 2. A sample of 330,069 UK Biobank participants with both scores on the reaction time test and genotyping data was used in this study.

### Genome-wide association analyses

Genotype–phenotype association analyses were performed within each cohort, using an additive model, on imputed SNP dosage scores. Adjustments for age, sex, and population stratification, if required, were included in the model. Cohort-specific covariates—for example, site or familial relationships—were also fitted as required. Cohort specific quality control procedures, imputation methods, and covariates are described in Supplementary Table S2. Quality control of the cohort-level summary statistics was performed using the EasyQC software^36^, which implemented the exclusion of SNPs with imputation quality < 0.6 and minor allele count < 25.

### Meta-analysis

A meta-analysis of the 56 CHARGE-COGENT cohorts was performed using the METAL package with an inverse variance weighted model implemented and single genomic control applied (http://www.sph.umich.edu/csg/abecasis/Metal). The two UK Biobank groups, VNR Assessment Centre and VNR Web-Based, were also meta-analysed using the same method. An inverse-variance weighted meta-analysis of the CHARGE-COGENT and UK Biobank summary results was then performed.

### Reaction Time Genome-wide association analysis

The GWAS of reaction time from the UK Biobank sample was performed using the BGENIE v 1.2 analysis package (https://jmarchini.org/bgenie/). A linear SNP association model was tested which accounted for genotype uncertainty. Reaction time was adjusted for the following covariates; age, sex, genotyping batch, genotyping array, assessment centre, and 40 principal components.

### Gene-based analysis (MAGMA)

Gene-based analysis was conducted using MAGMA^37^. All SNPs that were located within protein coding genes were used to derive a *P*-value describing the association found with general cognitive function and reaction time. The SNP-wise model from MAGMA was used and the NCBI build 37 was used to determine the location and boundaries of 18,199 autosomal genes. Linkage disequilibrium within and between each gene was gauged using the 1000 genomes phase 3 release^38^. A Bonferroni correction was applied to control for multiple testing; the genome-wide significance threshold was *P* < 2.75 × 10^−6^.

### Estimation of SNP-based heritability

Univariate GCTA-GREML analyses^39^ were used to estimate the proportion of variance explained by all common SNPs in four of the largest individual cohorts: ELSA, Understanding Society, UK Biobank, and Generation Scotland. Sample sizes for all of the GCTA analyses in these cohorts differed from the association analyses, because one individual was excluded from any pair of individuals who had an estimated coefficient of relatedness of > 0.025 to ensure that effects due to shared environment were not included. The same covariates were included in all GCTA-GREML analyses as for the SNP-based association analyses.

### Univariate Linkage Disequilibrium Score Regression (LDSC)

Univariate LDSC regression was performed on the summary statistics from the GWAS on general cognitive function and reaction time. The heritability Z-score provides a measure of the polygenic signal found in each data set. Values greater than 4 indicate that the data are suitable for use with bivariate LDSC regression^40^. The mean *χ*^2^ statistic indicates the inflation of the GWAS test statistics that, under the null hypothesis of no association (i.e. no inflation of test statistics), would be 1. For each GWAS, an LD regression was carried out by regressing the GWA test statistics (*χ*^2^) on to each SNP’s LD score (the sum of squared correlations between the minor allele frequency count of a SNP with the minor allele frequency count of every other SNP).

### Genetic correlations

Genetic correlations were estimated using two methods, bivariate GCTA-GREML^41^ and LDSC^40^. Bivariate GCTA was used to calculate genetic correlations between phenotypes and cohorts where the genotyping data were available. This method was used to calculate the genetic correlations between different cohorts for the general cognitive function phenotype. It was also employed to investigate the genetic contribution to the stability of UK Biobank’s participants’ verbal-numerical reasoning test scores in the assessment centre and then in web-based, online testing. In cases where only GWA summary results were available, LDSC was used to estimate genetic correlations between two traits—for example, general cognitive function and longevity—in order to estimate the degree of overlap between polygenic architecture of the traits. Genetic correlations were estimated between general cognitive function and reaction time and a number of health outcomes.

### Polygenic prediction

Polygenic profile score analysis was used to predict cognitive test performance in Generation Scotland, the English Longitudinal Study of Ageing, and Understanding Society. Polygenic profiles were created in PRSice^42^ using results of a general cognitive function meta-analysis that excluded the Generation Scotland, the English Longitudinal Study of Ageing, and Understanding Society cohorts. Polygenic profiles were also created based on the UK Biobank GWA reaction time results.

### Functional Annotation and Loci Discovery

Genomic risk loci were derived using FUnctional MApping and annotation of genetic associations (FUMA)^16^. Firstly, independent significant SNPs were identified using the SNP2GENE function and defined as SNPs with a *P*-value of ≤ 5 × 10^−8^ and independent of other genome wide significant SNPs at *R*^2^ < 0.6. Using these independent significant SNPs, candidate SNPs to be used in subsequent annotations were identified as all SNPs that had a MAF ≥ 0.0005 and were in LD of *R*^2^ ≥ 0.6 with at least one of the independent significant SNPs. These candidate SNPs included those from the 1000 genomes reference panel and need not have been included in the GWAS performed in the current study. Lead SNPs were also identified using the independent significant SNPs and were defined as those that were independent from each other at *R*^2^ < 0.1. Genomic risk loci that were 250kb or closer were merged into a single locus.

The lead SNPs and those in LD with the lead SNPs were then mapped to genes based on the functional consequences of genetic variation of the lead SNPs which was measured using ANNOVAR^43^ and the Ensembl genes build 85. Intergenic SNPs were mapped to the two closest up- and down-stream genes which can result in their being assigned to multiple genes. All SNPs found in 1000 genomes phase 3 were then annotated with a CADD score^44^, RegulomeDB score^45^, and 15-core chromatin states^46-48^.

The mapping of eQTLs was performed using each independent significant SNP and those in LD with it. eQTL information was obtained from the following databases: GTEx (http://www.gtexportal.org/home/), BRAINEAC (http://www.braineac.org/), Blood eQTL Browser (http://genenetwork.nl/bloodeqtlbrowser/), and BIOS QTL browser (http://genenetwork.nl/biosqtlbrowser/).

### Gene-set analysis

Gene-set analysis was conducted in MAGMA^37^ using competitive testing, which examines if genes within the gene set are more strongly associated with each of the cognitive phenotypes than other genes. Such competitive tests have been shown to control for Type 1 error rate as well as facilitating an understanding of the underlying biology of cognitive differences^49,50^. A total of 10 891 gene-sets (sourced from Gene Ontology^51^, Reactome^52^, and, SigDB^53^) were examined for enrichment of intelligence. A Bonferroni correction was applied to control for the multiple tests performed on the 10,891 gene sets available for analysis.

### Gene property analysis

In order to indicate the role of particular tissue types that influence differences in general cognitive function and reaction time, a gene property analysis was conducted using MAGMA. The goal of this analysis was to determine if, in 30 broad tissue types and 53 specific tissues, tissue-specific differential expression levels were predictive of the association of a gene with general cognitive function and reaction time. Tissue types were taken from the GTEx v6 RNA-seq database^54^ with expression values being log2 transformed with a pseudocount of 1 after winsorising at 50, with the average expression value being taken from each tissue. Multiple testing was controlled for using a Bonferroni correction.

